# Parafoveal vision reveals qualitative differences between FFA and PPA

**DOI:** 10.1101/2023.07.04.547646

**Authors:** Olga Kreichman, Sharon Gilaie-Dotan

**Affiliations:** 1 School of Optometry and Vision Science, Faculty of Life Science, Bar Ilan University, Ramat Gan, 5290002 Israel; The Gonda Multidisciplinary Brain Research Center, Bar Ilan University, Ramat Gan, 5290002 Israel; UCL Institute of Cognitive Neuroscience, London, WC1N 3AZ UK

**Keywords:** eccentricity effect, BOLD eccentricity effect, FFA PPA, faces, face inversion, house inversion, FMRI, visual cortex, parafovea

## Abstract

The center-periphery visual field axis guides early visual system organization with enhanced resources devoted to central vision leading to reduced peripheral performance relative to that of central vision (i.e., behavioral eccentricity effect) for most visual functions. The center-periphery organization extends to high-order visual cortex where for example the well-studied face-sensitive fusiform face area (FFA) shows sensitivity to central vision and place-sensitive parahippocampal place area (PPA) shows sensitivity to peripheral vision. As we have recently found that face perception is more sensitive to eccentricity than place perception, here we examined whether these behavioral findings reflect differences in FFA and PPA’s sensitivities to eccentricity. We assumed FFA would show higher sensitivity to eccentricity than PPA would, but that both regions’ modulation by eccentricity would be invariant to the viewed category. We parametrically investigated (fMRI, n=32) how FFA’s and PPA’s activations are modulated by eccentricity (≤8°) and category (upright/inverted faces/houses) while keeping stimulus size constant. As expected, FFA showed an overall higher sensitivity to eccentricity than PPA. However, both regions’ activation modulations by eccentricity were dependent on the viewed category. In FFA a reduction of activation with growing eccentricity (“BOLD eccentricity effect”) was found (with different amplitudes) for all categories. In PPA however, there were qualitative modulations of the BOLD eccentricity effect with mild BOLD eccentricity effect for houses but a reverse BOLD eccentricity effect for faces and no modulation for inverted faces. Our results emphasize that peripheral vision investigations are critical to further our understanding of visual processing both quantitatively and qualitatively.

**Significance statement:** Visual perception significantly deteriorates with growing distance from central vision (behavioral eccentricity effect) with varying degrees according to visual function. For some functions (but not others) deterioration follows the reduction of resources devoted to peripheral vision at early visual processing stages. While early stages of visual processing reflect retinal spatial arrangement, here we found that activations in high-order visual areas that are less sensitive to visual field location show a BOLD fMRI activation eccentricity effect that mirrors the behavioral eccentricity effect. Importantly, the BOLD eccentricity effects we found varied across regions and were modulated quantitatively and qualitatively by the viewed visual categories. Our results emphasize that peripheral vision investigations are critical to further our understanding of visual processing.

## Introduction

The visual system devotes extensive resources to central vision relative to peripheral vision and this center-periphery imbalance commences at the retina (Wässle et al., 1990) and continues into early visual cortex. The magnitude of this phenomenon was estimated in retinotopic area V1 and is termed the cortical magnification factor (CMF, Horton and Hoyt 1991).

Given these extensive resources dedicated to central vision, it is not surprising that many visual functions - from basic ones (Cowey and Rolls, 1974; Levi et al., 1985) to higher ones (Carrasco et al., 1995, 2003; Wolfe et al., 1998; Staugaard et al., 2016; Kreichman et al., 2020; Akselevich and Gilaie-Dotan, 2022) deteriorate significantly (performance-wise) as eccentricity (distance from central vision) increases (the behavioral eccentricity effect (Carrasco et al., 1995; Wolfe et al., 1998; Xue et al., 2023)). It was also shown in some cases that when the CMF is accounted for (such that peripheral stimuli are proportionally enlarged to elicit similar cortical surface activation as is elicited by central stimuli) peripheral performance can reach central performance (Cowey and Rolls, 1974; Carrasco and Frieder, 1997; Jigo et al., 2023).

However, in natural vision there is no compensation for CMF and an object in peripheral vision occupies the same retinal space as when it appears in central vision. Furthermore, since different visual functions do not deteriorate at a uniform rate (Kreichman et al., 2020; Akselevich and Gilaie-Dotan, 2022; Xue et al., 2023), additional task-, attention- or function-specific processes are likely to contribute to the eccentricity-based reductions. As we have recently found that face and house discrimination do not deteriorate similarly with eccentricity (Kreichman et al., 2020), here we assumed that this difference may be attributed to regions that are highly sensitive to these categories in high-order visual cortex.

Two of these regions are among the most investigated in high-order visual cortex: The “fusiform face area” (FFA, (Kanwisher et al., 1997; McCarthy et al., 1997; Grill-Spector and Malach, 2004; Gilaie-Dotan et al., 2008)) sensitive to faces and also known to show a foveal bias (Levy et al., 2001; Hasson et al., 2002), and the “parahippocampal place area” (PPA, (Epstein, Harris, Stanley, & Kanwisher, 1999; Epstein & Kanwisher, 1998)) sensitive to places and known to show preference to peripheral stimuli (Levy et al., 2001, 2004; Hasson et al., 2002; Malach et al., 2002). We were interested to parametrically assess FFA’s and PPA’s activation modulation by eccentricity for their preferred (faces in FFA, houses in PPA) and non-preferred (houses in FFA, faces in PPA) categories. We were also interested to examine how these regions’ activations are affected by eccentricity to inverted stimuli (Epstein et al., 2006; Kreichman et al., 2020).

To that end we ran an fMRI study (n=32) where we independently localized FFA and PPA and then tested how each of these region’s activation is affected by eccentricity in the parafovea (≤8°) for upright and inverted faces and houses when participants performed category-insensitive tasks (2 separate experiments with different cohorts of participants). To tap into the veridical sensitivity of these regions to eccentricity and parametrically examine eccentricity-based sensitivity across the visual field we kept stimulus size constant (not compensating for the CMF).

We assumed that both FFA and PPA would show eccentricity-based activation reductions (which we refer to here as the “BOLD eccentricity effect”) but each with a different rate (i.e., a different region-specific magnitude of BOLD eccentricity effect). Specifically, we hypothesized FFA would show a greater BOLD eccentricity effect (steeper reductions of activation with growing eccentricity) than PPA. In addition, we also assumed that each region would show its own “characteristic” BOLD eccentricity effect that would be category-independent given previous studies indicating (i) region-specific processing that shows category-dependency predominantly with respect to activation magnitude (Gilaie-Dotan et al., 2008; Grill-Spector et al., 2017b; Weiner et al., 2017) and (ii) that PPA showed a comparable peripheral bias for both faces and houses (Levy et al., 2001, 2004).

## Materials and Methods

### Participants

32 healthy participants (20 women, aged 18–43 years, mean age 28.6 years ± 5.5 SD, 29 right-handed, see more details in Table S1 at https://osf.io/m8czv/ (Kreichman and Gilaie-Dotan, n.d.)) participated in two fMRI studies (16 participants in upright and inverted ‘Count20’ experiments; 16 participants in upright ‘DBLstml’ experiment). Sample sizes were determined based on earlier studies investigating face-house sensitivities and eccentricity effects using similar cohort sizes (Levy et al., 2001: n1=13, n2=5, n3=6 participants; Hasson et al., 2002: n1=11, n2=6 participants; Levy et al., 2004: n1=11, n2=5, n3=8 participants, Kanwisher, Tong, & Nakayama, 1998: n1=20, n2=5, n3=5, Epstein & Kanwisher, 1998: n1=10, n2=6, n3=5). All participants reported having normal or corrected-to-normal vision and provided written informed consent to participate in the fMRI experiments before the experiments began. The Tel-Aviv Sourasky Medical Center ethics committee approved the experimental protocol.

### MRI setup

Participants were scanned in a Siemens Magnetom Prisma 3T scanner equipped with a standard 20-channel head coil at the Weizmann Institute of Science, Rehovot, Israel. Blood oxygenation level dependent (BOLD) contrast for the functional scans was obtained with gradient-echo echo-planar imaging (EPI) sequence (face/house experiments: TR=2500 ms, TE=30 ms, flip angle=80°, field of view (FoV)=216×216 mm^2^, matrix size=72×72, 40 axial slices of 3mm thickness (no gap) with an in-plane resolution of 3×3 mm^2^, covering the entire cortex; category localizer experiment: TR=3000 ms, TE=30 ms, flip angle=85°, FoV=210×210 mm^2^, matrix size=70×70, 48 axial slices of 3mm thickness (no gap) with an in-plane resolution of 3×3 mm^2^, covering the entire brain). A high-resolution whole brain anatomical T1-weighted magnetization prepared rapid acquisition gradient echo (MPRAGE) sequence was acquired for each participant (TR=2300 ms, TE=2.32 ms, flip angle=8°, FoV=240×240 mm^2^, 192 slices of 0.9 mm thickness with no gap, 1×1 mm^2^ in-plane resolution) to allow accurate cortical segmentation, reconstruction, and volume-based statistical analysis.

Stimuli were generated on a Windows 7 Enterprise PC and projected using an LCD projector onto a screen (31cm width X 35cm height) at the back end of the MRI tunnel with a viewing distance of 110 cm; scanner room was darkened during the experiments. Screen was viewed through a tilted mirror positioned over the participant’s forehead. Face/house experiments were run using an in-house developed platform for psychophysical and eye-tracking experiments (PSY) developed by Yoram S. Bonneh (Bonneh et al., 2015; Kreichman et al., 2020; Masarwa et al., 2022) running on a Windows PC and the category localizer experiment (Grossman et al., 2019) was run using "Presentation" software (Neurobehavioral Systems, Inc.). In addition, eye movements were recorded during face/house experiments using an “Eyelink−1000”, an MR-compatible eye-tracker (SR Research, Ontario, Canada) with a sampling rate of 500Hz equipped with a long-range lens. Eye movements of the right eye were recorded. For behavioral performance analyses button presses were recorded using a Fiber Optic Response Pad (fORP) 4-Button Curve Right response box (Current Designs, Philadelphia, USA).

### fMRI procedure and experimental design

Before beginning the experimental scans, each participant performed a standard 3 or 5-point eye-tracker calibration within the scanner (for 5 participants due to technical reasons we used previously stored calibration). The upright face/house experiment was always performed first, then the category localizer and anatomical scans (their order could be switched), and the inverted experimental runs were always performed last. All face/house experiments were block-design, each block presenting images of one category (face/house) and one eccentricity (0°, 4° or 8°). Participants were debriefed about the experiments after the MRI scan.

#### Main face/house experiment (upright ‘Count20’)

*Stimuli*. All face and house images were grayscale images presented on a black background. Face images subtending ∼1.9° x ∼2.3° (width x height) were full-front photographs of men with a neutral expression (taken from an earlier study (Gilaie-Dotan and Malach, 2007; Gilaie-Dotan et al., 2010) that modified images from 2 databases (CVL Face Database [http://www.lrv.fri.uni-lj.si/facedb.html]; AR Face Database [Martinez and Benavente 1998]). House images (as those used in Kreichman et al., 2020) were photographs of real houses and subtended ∼2° x ∼2.5° (width x height). Overall, we used 10 different face and 10 different house images. Due to screen limitations of the MRI setup face and house images centered at 8° on the vertical meridian (at the topmost and bottommost locations, see a video demonstration of the stimuli at https://osf.io/m8czv/) appeared cropped such that only the lower half of images presented at the upper part of the screen and only the upper half of the images presented at the lower part of the screen were visible (overall 6 out of 24 images during long blocks and 4 out of 16 images during short blocks of the 8° conditions appeared cropped; images at all other locations appeared in full size).

*Procedure*. Each run started with a 20 second fixation block followed by a block of 15 presented textures (200ms, 300ms ISI), after which another 7.5 seconds fixation block appeared and then the main face/house image conditions were presented. The run ended with a 12 second fixation block. The experiment was a 2 categories x 3 eccentricities block design experiment where each block included stimuli of one condition (one visual category (upright faces or houses) appearing at one eccentricity (center, 4° or 8°)). Each block type was repeated twice in each run (12 blocks per run) and block order was pseudo-random. 9 blocks presented 24 images and 3 pseudo-randomly chosen blocks presented 16 images. Images in each block were presented consecutively (200ms/image, 300ms ISI); in the central conditions, images appeared in the same location (screen center) and in the 4° and 8° conditions images appeared in 8 locations of the block’s eccentricity (in each quadrant and on the horizontal and vertical meridians, see Figure 1) in a pseudo-random order. Images could be repeated several times during a block and same images appeared across blocks. A white fixation circle was present throughout the duration of the run, at the end of each block the fixation circle turned green for 2500ms to indicate the block ended and response was expected. Between blocks, the white fixation circle continued to be present for 6 or 5 seconds (depending on whether it followed a long or a short block). Participants were instructed to fixate throughout the experiment, be aware of the surrounding area in which the targets could appear, and report at the end of each block (when the fixation circle turned green) whether there were more or less than 20 images displayed in that block (by pressing the right or the left buttons on the response box, respectively); no feedback was given (see Figure 1 for experimental paradigm and Figure 3 for behavioral performance). Each participant underwent two runs of the experiment, each run was of a different version of the experiment (with reversed block order between versions and different conditions chosen for the short blocks). Version order was counterbalanced across participants. Each run took 4 minutes 35 sec.

**Figure 1.**
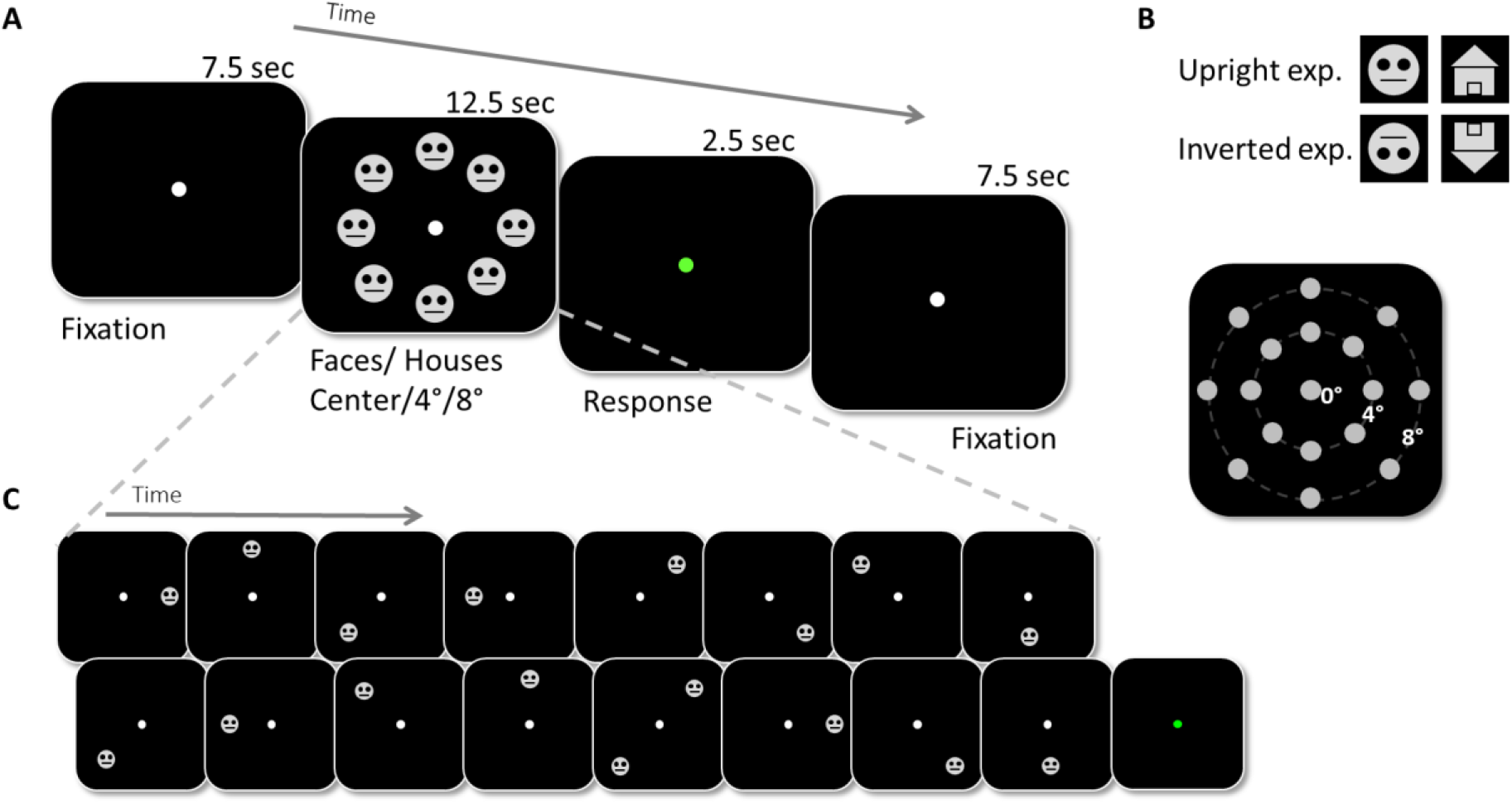
Experimental design of parafoveal face/house fMRI experiments. **(A)** Representative parafoveal upright face-block timeline. Each condition block (one category (faces or houses) at one eccentricity (0°, 4°, or 8°)) sequentially presented 16 or 24 images (200ms/image, 300ms ISI, white fixation present throughout) in pseudorandom order (see (C)), and ended with a green fixation (2.5s) indicating that response was expected (in ‘Count20’ to report whether more or less than 20 images appeared, in ‘DBLstml’ to report if 0-3 image pairs appeared, see Methods). Participants were instructed to keep fixation and be aware of the surrounding area in which the targets could appear. Each condition block was repeated twice in each experiment. **(B)** Representative stimuli for the upright and inverted ‘Count20’ and ‘DBLstml’ experiments and experimental stimuli locations (8 locations/parafoveal eccentricity). Inverted experiments had identical design with inverted stimuli. **(C)** Representative short block order (16 stimuli, ‘DBLstml’ experiment, expected response: 1 pair. Note that the experimental stimuli were real faces and of real houses and are fully described in the Methods section and the ones depicted here are only for illustration.

**Figure 2.**
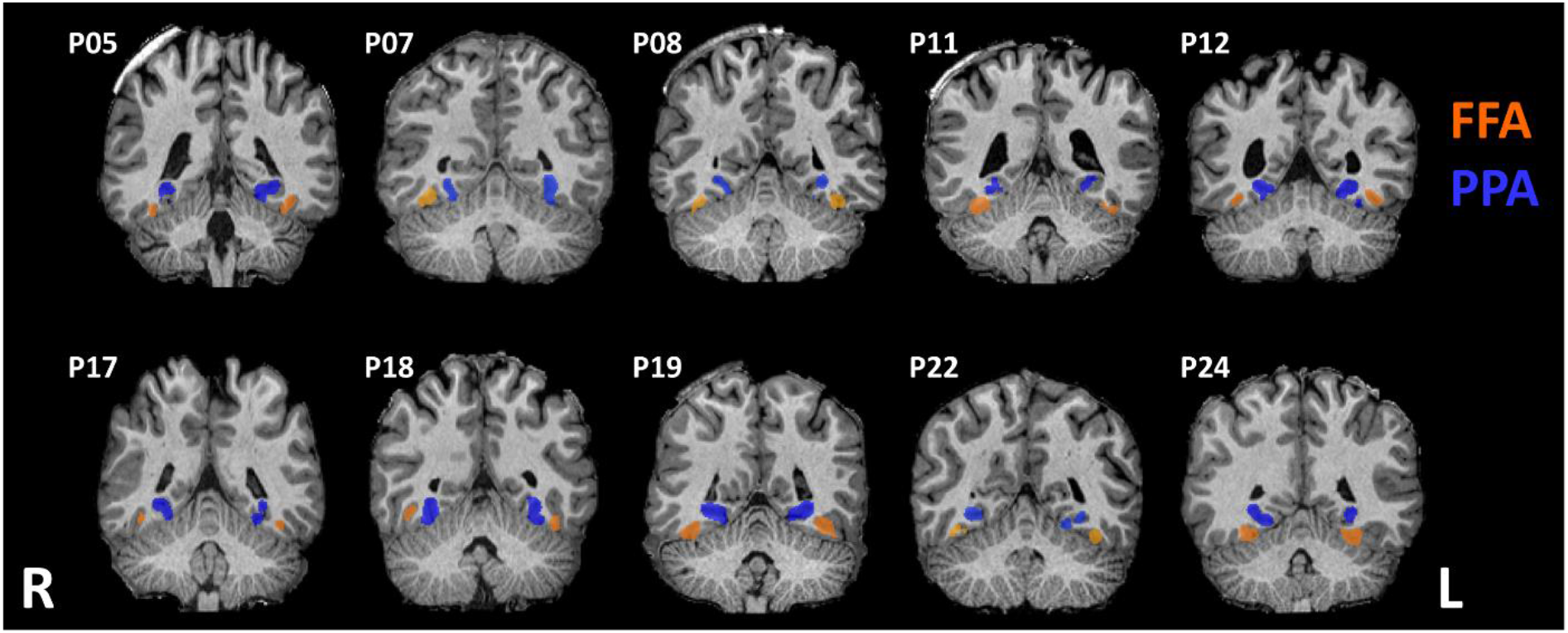
FFA and PPA independent localization (10 representative participants). Demonstration of individually defined FFA (in orange) and PPA (in blue) by the category localizer experiment (n=5 from ‘Count20’ experiment; n=5 from ‘DBLstml’ experiment; coronal views; see Methods). R/L – right/left hemisphere.

**Figure 3.**
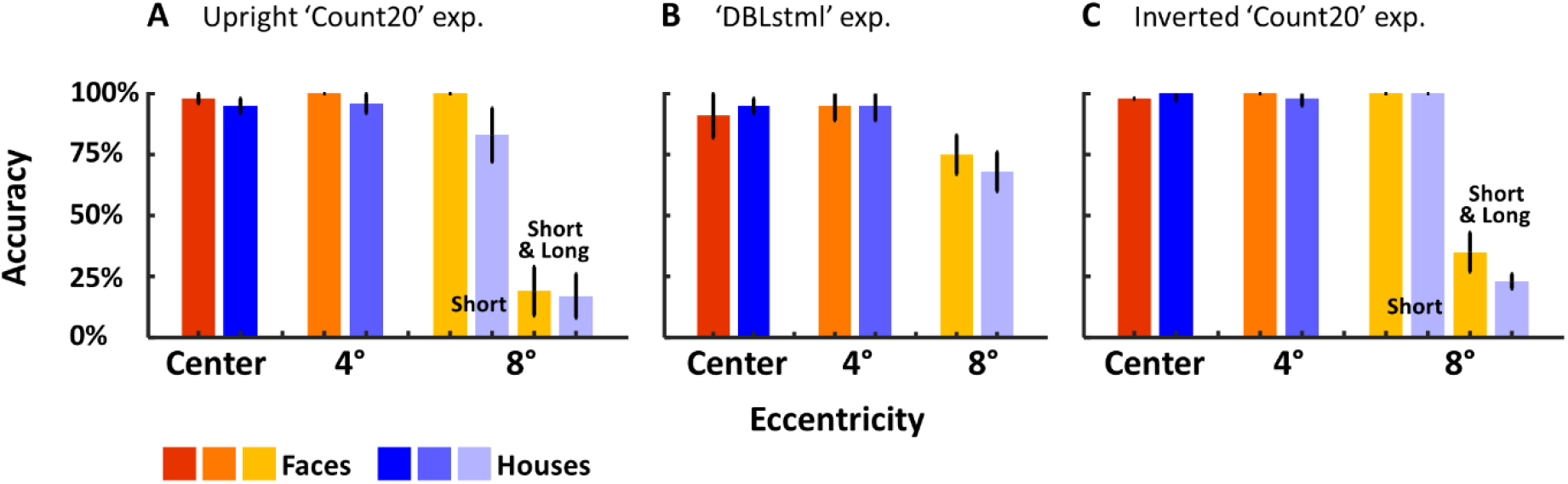
Behavioral performance by eccentricity and category for all experiments. Accuracy in **(A)** upright ‘Count20’ experiment (n=12), **(B)** ‘DBLstml’ experiment (n=11), and **(C)** inverted ‘Count20’ experiment (n=10). Face conditions in shades of red to yellow, house conditions in shades of blue. No differences between face and house accuracy were found across eccentricities and experiments. Behavior was almost at ceiling across eccentricities apart from the 8° long blocks (see Methods and Results). Error bars represent SEM.

#### Control face/house experiment 1 (upright ‘DBLstml’)

To control the possibility that the results of the upright ‘Count20’ experiment were task-dependent we ran an additional experiment with upright faces and houses on a different group of participants (‘DBLstml’ experiment). This experiment was identical in its design and stimuli to the upright ‘Count20’ experiment. Since we employed here a different task (see below), for several of the stimuli an additional same-category stimulus was added such that there were instances of 2 images appearing simultaneously on two opposing sides of fixation (see an example sequence in Figure 1 and a video demonstration of the stimuli at https://osf.io/m8czv/) and this double stimulus could happen once, twice, or three times in a block (double stimulus occurred in half of the blocks (6 of 12). The participants were not notified of the frequency of these pairs across the experiment and were instructed to fixate and be aware of the surrounding area in which the targets could appear, and report whether there were 0, 1, 2, or 3 simultaneous image pairs in a block (by pressing the corresponding button in the 4-buttons response box). This method of simultaneous image-pair presentation was also meant to facilitate fixation throughout the experimental session. There were two versions of the experiment differing in block order and number of paired-images stimuli in each block, each participant ran each version once. In the first version there were 4, 1 and 1 blocks with 1, 2 and 3 image pairs, respectively; in the second version there were 3, 2 and 1 blocks with 1, 2 and 3 image pairs, respectively. Images used in this experiment were same as those used in the ‘Count20’ experiment. Each run took 4min and 35 sec.

#### Control face/house experiment 2 (inverted ‘Count20’)

Experimental design was identical to that used in the upright ‘Count20’ paradigm except that stimuli were inverted.

#### Category localizer experiment

Each participant underwent a blocked design visual category localizer experiment (Gilaie-Dotan et al., 2008, 2009, 2010, 2013; Berkovich-Ohana et al., 2020) to identify the cortical regions preferentially activated by specific visual categories (faces, houses, objects, body parts, or patterns (Grossman et al., 2019)). Each block lasted 9s and included 9 images (6°×6°, 800ms/image, 200ms ISI) that were all from the same visual category presented on a gray background. Each block was followed by a 6 second blank screen. Blocks of each category were repeated 7 times in a pseudorandom order across the experiment. The experiment lasted 550 seconds. The first and last blocks of the experiment were fixation blocks that lasted 21 seconds and 9 seconds, respectively. A central fixation point was present throughout the experiment. Participants were instructed to perform a 1-back memory task and report via button presses whether the presented stimulus was identical to or different than the previous stimulus. Image repetition occurred once or twice in each block.

#### fMRI data preprocessing and analysis

fMRI data were analyzed with the BrainVoyager software version 21.4 (Brain Innovation, Maastricht, The Netherlands). The first 2 TRs (volumes) of each functional scan were discarded. Preprocessing of functional scans included 3D motion correction, slice scan time correction, linear trend removal and high-pass filtering (filtering out frequencies lower than 3 cycles across the experiment). Functional data were incorporated into 3D normalized MNI space where all further analyses were performed.

Exclusions performed on the data per participant (see https://osf.io/m8czv/): runs with head motion larger than 1mm in 1 of the 6 motion directions in both runs were excluded from the analyses (participants with one run excluded based on head motion: upright ‘Count20’: n=1, inverted Count20: n=1, ‘DBLstml’: n=2) and participants whose both upright runs were excluded based on head motion were discarded from the study (upright ‘Count20’: n=2; ‘DBLstml’: n=1). On top of this, 3 additional participants had both of their inverted ‘Count20’ runs excluded based on head motion (so their upright runs but not their inverted runs were included). 4 additional participants only underwent 1 run in the upright ‘DBLstml’ experiment. In addition, 1 participant was excluded due to chance level behavioral results during central blocks of upright ‘DBLstml’ experiment. An additional participant’s data of one run of upright ‘Count20’ exp. could not be analyzed due to technical issues.

#### ROI Analysis

Functional regions of interest (ROIs) in the right and left hemispheres were defined individually for each participant using the category localizer experiment (Gilaie-Dotan et al., 2008, 2009, 2010). Face-sensitive area in the lateral fusiform gyrus (“FFA” Kanwisher et al., 1997; McCarthy et al., 1997) was defined by preferential activation to faces relative to houses (contrast: faces > houses, p<0.003 at the voxel level, uncorrected), and place-sensitive area in the collateral sulcus (CoS) and parahippocampal gyrus (“PPA” (Epstein and Kanwisher, 1998)) was defined by preferential activation to houses relative to faces (contrast: houses > faces, p<0.003 at the voxel level, uncorrected), see Figure 2. In some participants, some ROIs could not be delineated in one of the hemispheres since they did not show the expected category selectivity (see Table S2 at https://osf.io/m8czv/ for a detailed list). For 2 participants with missing category localizer experimental data (1 was excluded due to head movements, 1 did not complete the localizer experiment) the localization of face and place related regions were based on contrasting the central face and central house blocks of our main fMRI experiment (upright face/house experiment; FFA by faceCenter > houseCenter, PPA by houseCenter > faceCenter) in combination with the expected anatomical locations. Table 1 presents average MNI coordinates and size (number of voxels) across participants for each ROI (see Table S2 at https://osf.io/m8czv/ for individual participants’ ROI MNI coordinates).

**Table 1.** Independently defined FFA and PPA ROI details for each of the upright experiments. For both the ‘Count20’ experimental cohort (n=14, top) and the ‘DBLstml’ cohort (n=14, bottom) MNI coordinates and number of voxels are presented for each ROI (average and standard error across participants). n represents number of participants that had each ROI defined after exclusion; in parentheses is the number of participants that also underwent the Inverted Count20 experiment (note that the group mean ROI coordinates presented here were defined solely based on the upright experiment cohort). R/L – right/left hemisphere (for a detailed list by participant see https://osf.io/m8czv/).

|  |  |  | Average |  |  |  | SE |  |  |  | n |
| --- | --- | --- | --- | --- | --- | --- | --- | --- | --- | --- | --- |
|  |  |  | X | Y | Z | Voxel count | X | Y | Z | Voxel count |  |
| Participants from 'Count20' exp. (n=14) | FFA | L | -41.08 | -51.78 | -19.44 | 431.25 | 1.02 | 1.50 | 1.14 | 73.93 | 12 (10) |
|  |  | R | 41.94 | -53.10 | -17.32 | 713.92 | 0.82 | 1.52 | 0.99 | 90.61 | 13 (10) |
|  | PPA | L | -28.62 | -50.86 | -10.37 | 1513.84 | 0.48 | 0.94 | 0.58 | 243.59 | 13 (10) |
|  |  | R | 29.64 | -50.10 | -10.08 | 1703.07 | 0.77 | 1.06 | 0.67 | 352.25 | 14 (11) |
| Participants from 'DBLstml' exp. (n=14) | FFA | L | -38.90 | -50.81 | -19.04 | 612.25 | 1.02 | 1.48 | 0.63 | 88.17 | 12 |
|  |  | R | 39.94 | -50.96 | -17.41 | 602.42 | 0.93 | 1.18 | 0.83 | 97.17 | 14 |
|  | PPA | L | -27.45 | -50.05 | -10.54 | 1115.21 | 0.61 | 0.73 | 0.73 | 141.25 | 14 |
|  |  | R | 27.87 | -48.74 | -9.08 | 1350.50 | 0.48 | 0.64 | 0.61 | 169.07 | 14 |

For each participant, event-related time course of activation was based on both runs each participant underwent (two versions of the experiment). For each participant and each ROI (FFA-R, FFA-L, PPA-R, PPA-L) we sampled the average event-related time course (upright ‘Count20’: Fig. 4A, inverted ‘Count20’ and ‘DBLstml’: Fig. 5A and D). The peak response was defined as the average response during TRs 3 and 4. The normalized response was defined as the response of each condition divided by the response of the ROI’s preferred category at 0° (Figure 4B and 5B). The slope of the response in each ROI was defined as the change in peak response by eccentricity between central 0° to 8° and this was calculated for the preferred and for the non-preferred categories separately (Figure 4C, 5C and 5F). Eccentricity-specific category selectivity was calculated by subtracting the response of the non-preferred category from that of the preferred category at each eccentricity (i.e., for FFA: faceCenter vs houseCenter, face-4° vs house-4°, face-8° vs house-8°; for PPA: face conditions were subtracted from house conditions, Fig. 7).

**Figure 4.**
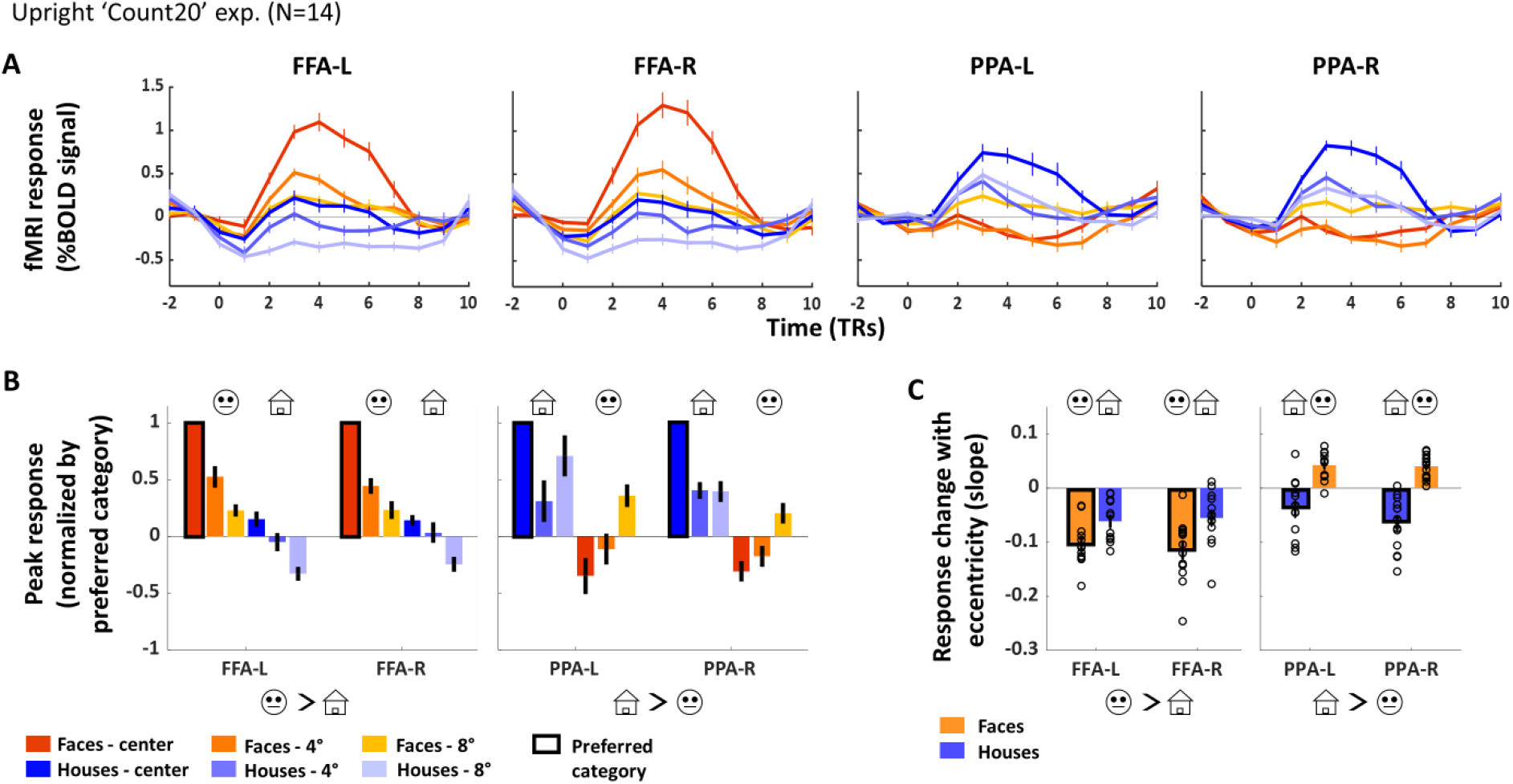
FFA and PPA activation modulation in the parafovea to preferred and non-preferred categories (upright ‘Count20’ experiment, n=14). **(A)** Mean activation (% signal change, y axis) for each experimental condition as a function of time (by TRs on the x axis) in each ROI across participants (FFA-L (n=12), FFA-R (n=13), PPA-L (n=13), PPA-R (n=14)), faces in dark to light orange with growing eccentricity, houses in dark to light blue with growing eccentricity. Category selectivity in central vision is evident in all ROIs by higher activation to the central preferred vs central non-preferred category. **(B)** Peak responses (mean of 3^rd^ and 4^th^ TRs from A) of each condition relative to that of central preferred category were calculated (normalized response); black borders demarcate each ROI’s activation to it central-preferred category. **(C)** Response modulation with eccentricity (slope from 0° to 8°, see Methods) for preferred and non-preferred categories for each ROI; a negative slope represents the BOLD eccentricity effect (activation reduction with growing eccentricity). Dots represent individual data. Note that FFA showed a BOLD eccentricity effect for both categories (albeit to different extents), but PPA showed qualitatively different modulations with a typical BOLD eccentricity effect for its preferred category but a reverse effect (positive slope reflecting a reverse/negative BOLD eccentricity effect) for the non-preferred category. Error bars represent SEM across participants.

**Figure 5.**
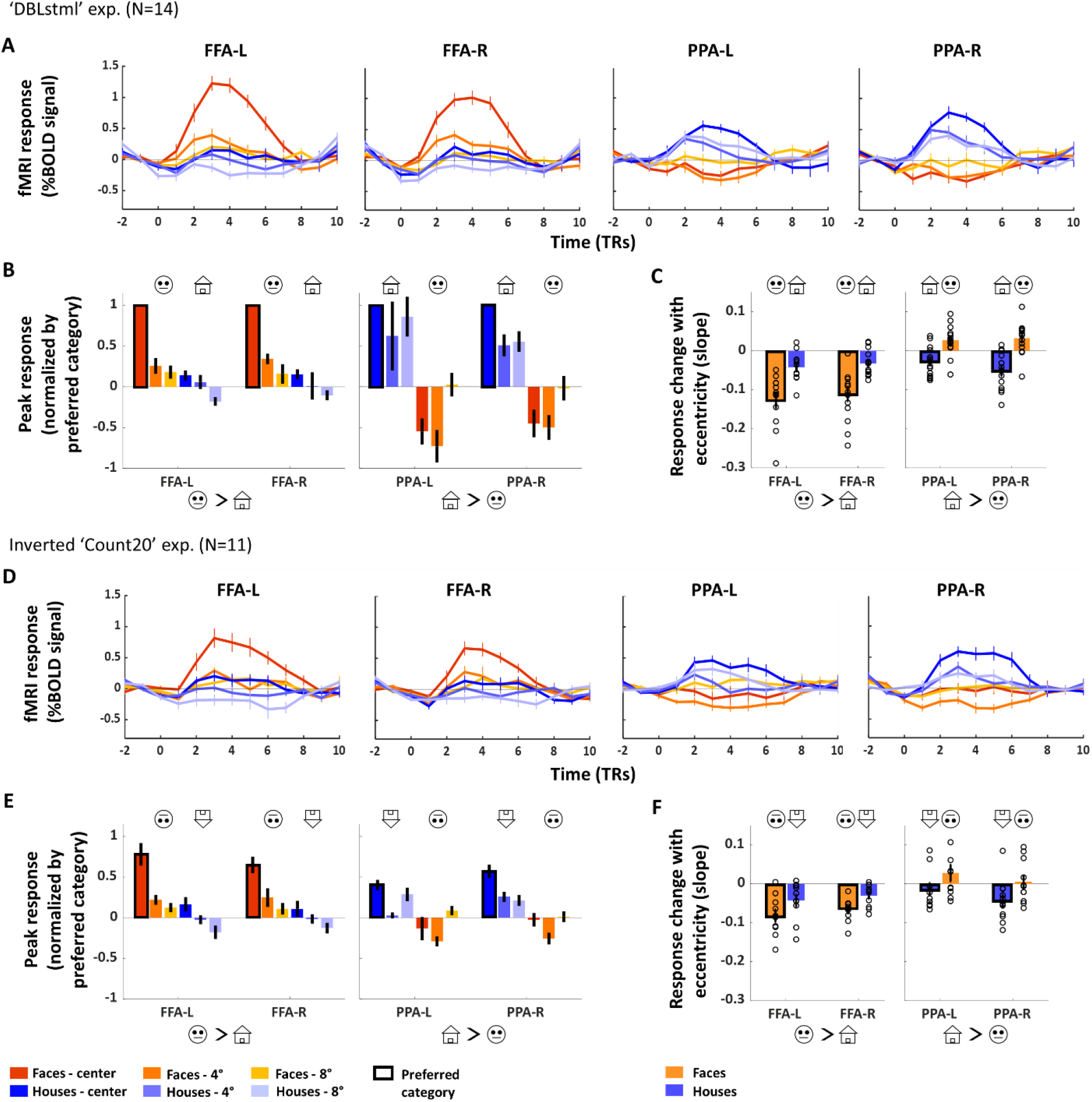
FFA and PPA activation modulation in the parafovea to preferred and non-preferred categories in control experiments (‘DBLstml’, n=14; inverted ‘Count20’, n=11). Notations as in Figure 3 with **(A-C) for** ‘DBLstml’ results (FFA-L (n=12), FFA-R (n=14), PPA-L (n=14), PPA-R (n=14)) and **(D-F)** for inverted ‘Count20’ (FFA-L (n=10), FFA-R (n=10), PPA-L (n=10), PPA-R (n=11)). Since inverted stimuli are not considered optimal for FFA and PPA, in **(E)** peak responses are not normalized. Note that in (C) similar to the ‘Count20’ results (Figure 4), in the ‘DBLstml’ experiment - while FFA showed the BOLD eccentricity effect (evident by negative slopes) for both preferred and non-preferred categories, PPA showed quantitatively different modulations by viewed category (negative slope for its preferred (houses) category vs a positive slope to its non-preferred (faces) category (see Results)). Also, in the inverted experiment we found that while FFA showed similar effects as for upright stimuli (F), for PPA there was no modulation by eccentricity to inverted faces. Error bars represent SEM.

#### Fixation assessment

Since our experiment aimed to evaluate activation to peripheral stimuli, eye movements were recorded to validate fixation was kept throughout the experimental sessions. Fixation analysis was performed offline. To evaluate participant’s fixation performance, for each block, we calculated the percentage of time that the participant’s gaze was within 1.5° from the fixation point (inter-stimulus intervals were not included in this analysis). We then computed a fixation score for each eccentricity by averaging across 4 blocks of each eccentricity (taking into account blocks of faces and of houses together), this was done separately for each run. We defined good fixators as participants that kept fixation at least 80% of the time during the 8° blocks. We evaluated the effect of fixation on activation by comparing activation of participants with good vs. poor fixations (see Tables S1 and S3 at https://osf.io/m8czv/).

#### Behavioral performance analysis

During each experiment participants were asked to keep fixation throughout the blocks and perform the behavioral task while being aware of the peripheral area in which stimuli could appear. Both behavioral tasks were aimed to keep participants attentive during the experiments while not demanding category-specific attention.

The partial visibility of the 8°images on the vertical meridian (extending beyond the visible visual display, see ‘Main face/house experiment (upright ‘Count20’)’ in Methods) may have influenced performance in the long blocks of the ‘Count20’ experiment, so we mostly relied on the short blocks’ performance for 8° blocks.

In the upright ‘Count20’, ‘DBLstml’ and inverted ‘Count20’ experiments there were 2, 3 and 1 participants (respectively) whose responses were not recorded due to technical issues resulting in behavioral results reported for n=12 (upright ‘Count20’), n=11 (‘DBLstml’) and n=10 (inverted ‘Count20’); see Figure 3 and Table 2. Note that one participant in the upright ‘Count20’ experiment did not respond to the central conditions, and for one participant in the upright ‘Count20’ exp. and 2 participants in the ‘DBLstml’ exp. only seven out of 8 repetitions were available for the central condition. See https://osf.io/m8czv/ for more details.

**Table 2.** Group behavioral performance by eccentricity and category for upright ‘Count20’, ‘DBLstml’ and inverted ‘Count20’ experiments. Average accuracy (% correct) ± SEM are presented. See Figure 3, ‘Behavioral performance analysis’ in Methods and https://osf.io/m8czv/ for further details.

|  |  |  | Upright 'Count20'<br>(n=12) | 'DBLstml'<br>(n = 11) | Inverted 'Count20'<br>(n=10) |
| --- | --- | --- | --- | --- | --- |
| Center | Faces |  | 97.7 ± 2.2 | 90.9 ± 9.1 | 97.5 ± 2.5 |
|  | Houses |  | 95.5 ± 2.9 | 95.5 ± 3 | 100 ± 0 |
| 4° | Faces |  | 100 ± 0 | 95.5 ± 6.2 | 100 ± 0 |
|  | Houses |  | 95.8 ± 4.2 | 95.5 ± 6.2 | 97.5 ± 2.5 |
| 8° | Short blocks | Faces | 100 ± 0 |  | 100 ± 0 |
|  |  | Houses | 83.3 ± 11.2 |  | 100 ± 0 |
|  | Short+long blocks | Faces | 37.5 ± 7.8 | 75.0 ± 7.5 | 35.0 ± 7.6 |
|  |  | Houses | 33.3 ± 7.7 | 68.2 ± 7.6 | 22.5 ± 2.5 |

### Statistical analyses

Statistical analyses (ANOVA, post-hoc and Wilcoxon signed-rank test) were performed using R studio (version 2021.9.0.351, RStudio Team, 2020). Repeated measures ANOVAs were performed using the R get_anova_table() function that automatically apply the Greenhouse-Geisser sphericity correction only to factors violating the sphericity assumption (i.e., Mauchly’s test significant p-value ≤ 0.05). One-sample, 2-sided Wilcoxon signed rank tests were always preformed relative to the zero vector.

## Results

### Behavioral performance

We deliberately chose a behavioral task that would near ceiling performance so that task difficulty would be comparable across eccentricities and visual categories and thus will allow investigating FFA and PPA’s sensitivity to visual categories regardless of task modulations. Accuracy for the central and 4° conditions was close to ceiling in the upright experiments (‘Count20’, ‘DBLstml’) and slightly lower for the inverted ‘Count20’ experiment (Figure 3 and Table 2). In the 8° condition (‘Count20’ experiment, upright and inverted) we found that performance for the short blocks alone was high (as for central and 4° blocks), but for the long and short blocks combined it was close to chance (can possibly be attributed to the partial visibility of the 8° stimuli on the vertical meridian, Figure 3 and Table 2). Importantly, performance appeared to be comparable for the face and house conditions at all eccentricities.

### FFA’s and PPA’s activation modulation by eccentricity

#### Main experiment – upright ‘Count20’ experiment

FFA and PPA were independently localized using an external localizer (see Figure 2) with anatomical locations (face selective areas in the lateral fusiform and house-selective areas in the collateral sulcus) and coordinates (see Table 1 and Table S2 at https://osf.io/m8czv/) comparable to those found in earlier studies (e.g. (Ishai et al., 1999; Hasson et al., 2002)). Afterwards, the experimental time courses (upright ‘Count20’ experiment, n=14) of each of our 4 ROIs (FFA-R/L, PPA-R/L) were analyzed. We first verified that these ROIs showed the expected category selectivity for the central experimental stimuli in our experiment as expected from earlier studies (e.g. Gilaie-Dotan et al., 2008) and indeed this is what we found for each of the ROIs (paired, 2-sided Wilcoxon signed rank test: right/left FFA: faceCenter > houseCenter: p’s < 10^-3^, in right/left PPA: houseCenter > faceCenter: p’s < 10^-3^, see and Figures 4 and 5, Table 3). Furthermore, as can be seen in Figures 4A and 5A, activation levels to the preferred categories of each of the ROIs (for faces in FFA-R and FFA-L, for places in PPA-R and PPA-L) appeared to decrease with eccentricity as would be expected from the behavioral eccentricity effect (Carrasco et al., 1995; Wolfe et al., 1998).

**Table 3.**
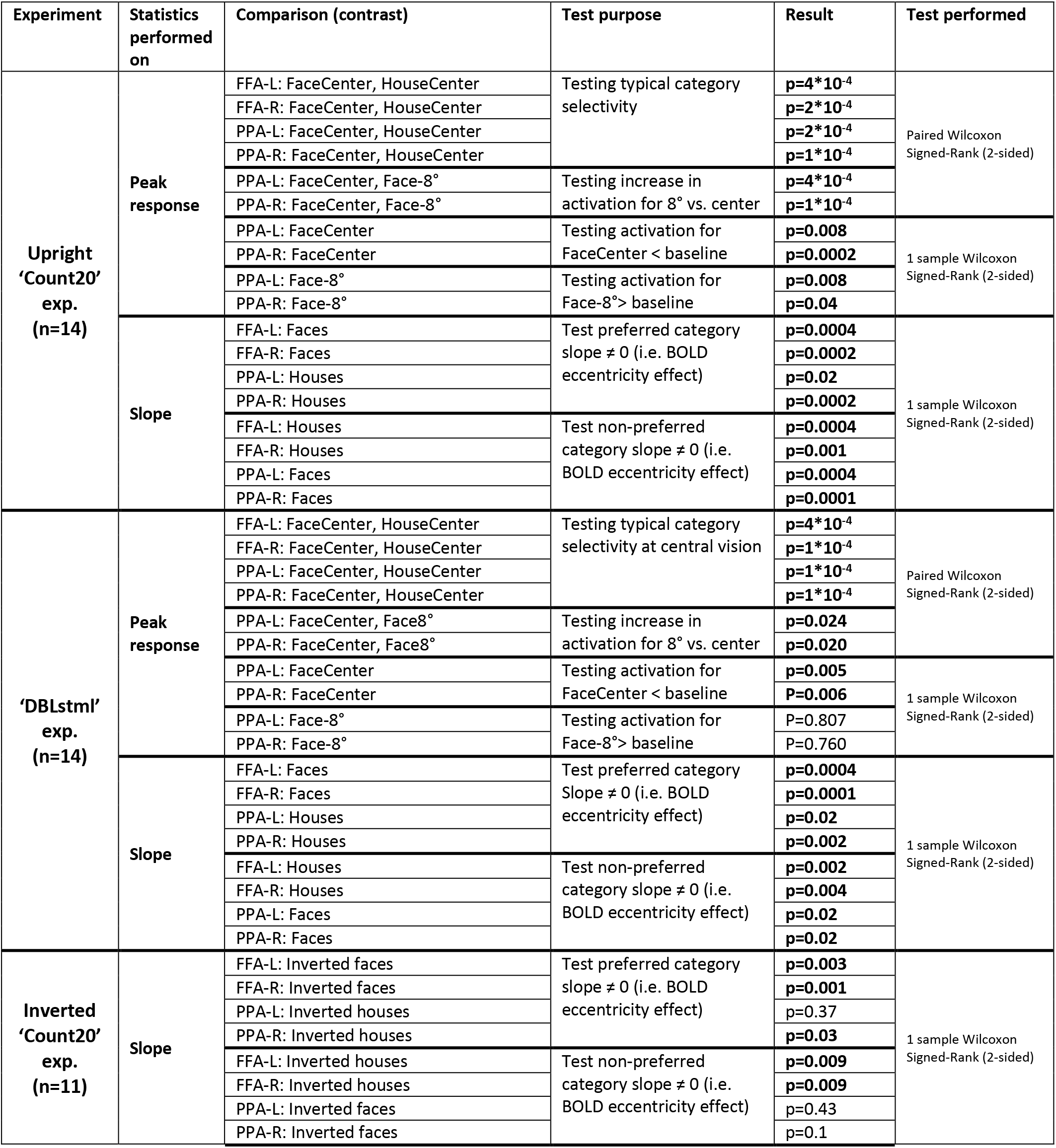
Complementary statistical comparisons. For each statistical contrast its full details are provided and the outcome in Result column. Significant results in bold. See Results for specific details.

To examine whether these apparent BOLD eccentricity effects were significant for each ROI, whether any BOLD eccentricity effects were modulated by stimulus category (preferred vs non-preferred stimuli), and whether there were any right-left hemispheric differences for FFA and/or PPA, we subjected the event-related peak activations (see Figure 4A and 5A and Methods) in right and left FFA and in right and left PPA to 3-way ANOVAs (eccentricity _(0°, 4°, 8°)_ x category _(preferred/non-preferred)_ x hemisphere _(R/L hemisphere)_). As expected, for both FFA and for PPA we found a significant BOLD eccentricity effect (all p’s < 0.01; Table 4), a significant stimulus category effect (all p’s < 10^-5^), and importantly, an interaction between category and eccentricity (FFA: p=6*10^-4^, PPA: p=1.3*10^-6^). This indicates that for both FFA and PPA their eccentricity-based activation modulations were dependent on their stimulus category preference such that the modulations for their preferred and non-preferred categories were different). No differences between right and left hemispheres were found, however a significant 3-way interaction (hemisphere X category X eccentricity) was found in PPA (F(2,24)=5.82 p = 0.009, Table 4).

**Table 4.** Statistical examination of hemisphere, category and eccentricity effect for FFA and PPA in each experiment. For each ROI and each experiment, the results of a repeated measures 3-way ANOVA on peak response with hemisphere (R, L), category-preference, and eccentricity as factors are presented (see Results for further details). Pref - preferred category, Non-pref - non-preferred category, R/L – right/left hemisphere. Significant results in bold.

| Experiment | ROI | Factors | Main effects | Interactions |
| --- | --- | --- | --- | --- |
| 'Count20' exp. | FFA (n=11) | Hemisphere R, L | F(1,10)=0.17, p=0.68 | Hemisphere X Category<br>F(1,10)=0.27, p=0.61<br>Hemisphere X Eccentricity<br>F(2,20)=0.41, p=0.66<br>Category X Eccentricity<br>F(2,20)=10.78, **p=6\*10<sup>-4</sup>**<br>Hemisphere X Category X Eccentricity<br>F(1.2,12.01)=0.43, p=0.56 |
|  |  | Category Pref, Non-pref | F(1,10)=140.86, **p=3\*10<sup>-7</sup>** |  |
|  |  | Eccentricity 0°, 4°, 8° | F(2,20)=48.43, **p=2\*10<sup>-8</sup>** |  |
|  | PPA (n=13) | Hemisphere R, L | F(1,12)=0.88, p=0.36 | Hemisphere X Category<br>F(1,12)=0.43, p=0.52<br>Hemisphere X Eccentricity<br>F(2,24)=2.9, p=0.07<br>Category X Eccentricity<br>F(2,24)=25.05, **p=1\*10<sup>-6</sup>**<br>Hemisphere X Category X Eccentricity<br>F(2,24)=5.82, **p=0.009** |
|  |  | Category Pref, Non-pref | F(1,12)=72.94, **p=1\*10<sup>-6</sup>** |  |
|  |  | Eccentricity 0°, 4°, 8° | F(2,24)=7.84, **p=0.003** |  |
| 'DBLstml' exp. | FFA (n=12) | Hemisphere R, L | F(1,11)=2.91, p=0.1 | Hemisphere X Category<br>F(1,11)=6.1, **p=0.03**<br>Hemisphere X Eccentricity<br>F(2,22)=0.83, p=0.4<br>Category X Eccentricity<br>F(2,22)=9.43, **p=0.003**<br>Hemisphere X Category X Eccentricity<br>F(2,22)=3.49, **p=0.04** |
|  |  | Category Pref, Non-pref | F(1,11)=57.189, **p=1\*10<sup>-5</sup>** |  |
|  |  | Eccentricity 0°, 4°, 8° | F(2,22)=32.8, **p=2\*10<sup>-7</sup>** |  |
|  | PPA (n=14) | Hemisphere R, L | F(1,13)=4.35, p=0.057 | Hemisphere X Category<br>F(1,13)=4.94, **p=0.04**<br>Hemisphere X Eccentricity<br>F(2,26)=2.04, p=0.1<br>Category X Eccentricity<br>F(2,26)=5.53, **p=0.02**<br>Hemisphere X Category X Eccentricity<br>F(2,26)=2.41, p=0.1 |
|  |  | Category Pref, Non-pref | F(1,13)=45.81, **p=1\*10<sup>-5</sup>** |  |
|  |  | Eccentricity 0°, 4°, 8° | F(2,26)=2.94, p=0.07 |  |

To check whether our results were potentially affected by poor fixations we also compared these brain activations between participants with good vs those with poor fixations for each of the ROIs separately. While only half of the participants in the upright ‘Count20’ experiment (7 of 14) were classified as having good fixation behavior (see Methods), we found that activation levels were not affected by fixation quality in any of the ROIs (3-way mixed-design ANOVAs with fixation quality _(good/poor)_ X eccentricity _(0°, 4°, 8°)_ X category _(preferred/non-preferred)_ on peak responses revealed no significant effect of fixation quality and no significant interaction between fixation quality and condition in all ROIs; see Table S3 at https://osf.io/m8czv/). Therefore, no participants were excluded from further analyses.

We normalized the responses within each ROI relative to the activation to its preferred category at central vision (see Methods and Figures 4B). In FFA, as can be seen in Figure 4B, we found a more gradual decrease in activation with growing eccentricity for its preferred (faces) than for its non-preferred (houses) category such that the activation to central and parafoveal faces remained above baseline (0° to 8° activity reduction for faces: FFA-L: 76%±18%(SD), FFA-R: 76%±28%(SD)) whereas activation to parafoveal houses at 8° declined to below baseline activity. In PPA, the decrease in activation with eccentricity for its preferred category (from 0° to 8° for houses: PPA-L: 28%±65%(SD), PPA-R: 60%±34%(SD)) appeared to be more moderate than that found for faces in the FFA (∼76% decrease), as we anticipated. However, more importantly, in PPA we observed that for its non-preferred category (faces) there was a substantial deviation from the anticipated BOLD eccentricity effect. Specifically, while the typical BOLD eccentricity effect is evident by reduced activation with growing eccentricity, in PPA we found an unexpected *increase* (rather than decrease) of activation with growing eccentricity for the non-preferred category (faces) such that faces at 8° significantly activated PPA above baseline (one-sample, 2-sided Wilcoxon signed rank test: face-8°: PPA-L: p=0.008, PPA-R: p=0.04, see Table 3) while central faces significantly deactivated it (one-sample, 2-sided Wilcoxon signed rank test: faceCenter: PPA-L: p=0.008, PPA-R: p=0.0002, see Table 4). This eccentricity-based rise of activity was significant (paired, 2-sided Wilcoxon signed rank test: faceCenter vs face-8°: PPA-L p=0.0004; PPA-R p=0.0001, see Figure 4B and Table 3).

To estimate and quantify the eccentricity-related activation modulations we defined an index that reflected the slope of response change with eccentricity (from 0° to 8°) for each ROI, for each category (see Figure 4C and https://osf.io/m8czv/ Table S4, (Kreichman et al., 2020)). For their preferred categories both PPA and FFA showed an anticipated BOLD eccentricity effect [significant negative slopes with one-sample, 2-sided Wilcoxon signed rank test: houses in PPA (PPA-L: p=0.02, PPA-R: p=0.0002), faces in FFA (FFA-L: p=0.0004, FFA-R: p=0.0002); see Fig. 4C] where PPA showed a quantitatively smaller BOLD eccentricity effect (smaller slope) than that found in the FFA [repeated 2-way ANOVA with ROI _(FFA/PPA)_ X hemisphere _(R/L)_ on the preferred category slopes revealed a significant ROI (but not hemisphere) effect (p=0.02); see Table 5]. Importantly, for its non-preferred category PPA showed an unexpected *reverse* eccentricity effect (enhancement of activation with growing eccentricity). This reverse BOLD eccentricity effect for PPA’s non-preferred category (faces) was qualitatively different than that found for its preferred category [(significantly negative slope for houses PPA-L: p=0.0002, PPA-R: p=0.02, one-sample 2-sided Wilcoxon signed rank test) but significantly positive slope for faces (PPA-L: p=0.0004, PPA-R: p=0.0001), see Table 3 and Figure 4C] and from that found in the FFA for the non-preferred category (houses) where slopes were significantly negative in line with the anticipated BOLD eccentricity effect (significantly negative slope for houses: FFA-L: p=0.0004, FFA-R: p=0.001, one-sample 2-sided Wilcoxon signed rank test; see Table 3).

**Table 5.** FFA vs PPA preferred category slope differences. For each upright experiment the results of a 2-way repeated measures ANOVA (with ROI and hemisphere as factors) on the preferred category slope are presented (see Results). Significant results in bold. Note that in both experiments a significant difference between FFA and PPA was found in their preferred category slope reflecting FFA’s higher sensitivity to eccentricity.

| fMRI experiment | Factor | Main effect | Interactions |
| --- | --- | --- | --- |
| **Upright ‘Count20’**<br>**exp.**<br>**(n=10)** | ROI<br>FFA, PPA | **F(1,9)=6.9, p=0.02** | F(1,9)=1.61, p=0.23 |
|  | Hemisphere<br>R, L | F(1,9)=1.28, p=0.28 |  |
| **‘DBLstml’**<br>**exp.**<br>**(n=12)** | ROI<br>FFA, PPA | **F(1,11)=16.7, p=0.002** | F(1,11)=4.77, p=0.051 |
|  | Hemisphere<br>R, L | F(1,11)=0.38, p=0.54 |  |

In summary, in the upright ‘Count20’ experiment we found, in line with our hypothesis (i) that the expected face-house selectivity in each of the ROIs, (ii) that FFA declined more rapidly with eccentricity than PPA did, and in contrast to our hypothesis that (iii) in both regions the eccentricity-based decline was modulated by visual category (see Table 6), and importantly (iv) PPA showed a qualitative difference in its decline such that for houses the BOLD eccentricity effect was observed but for faces it showed a negative/reverse BOLD eccentricity effect (growing activation with eccentricity).

**Table 6.** Effects of category preference on slope for each region, by experiment. For each ROI repeated 2-way ANOVAs with category preference X hemisphere on the slope (change of peak response between central to 8° stimuli, see Methods). Significant effects are indicated in bold. Note that for both regions there is a significant effect of category preference on the slope in the upright experiments. Pref/Non-pref indicate for each region it preferred and non-preferred categories.

| fMRI experiment |  | Factor | Main effect | Interactions |
| --- | --- | --- | --- | --- |
| Upright 'Count20' exp. | FFA (n=11) | Category Pref, Non-pref | F(1,10)=17.47, **p=0.002** | F(1,10)=1.15, p=0.30 |
|  |  | Hemisphere R, L | F(1,10)=0.79, p=0.39 |  |
|  | PPA (n=13) | ROI Pref, Non-pref | F(1,12)=53.12, **p=9\*10<sup>-6</sup>** | F(1,12)=8.63, **p=0.01** |
|  |  | Hemisphere R, L | F(1,12)=3.18, p=0.1 |  |
| 'DBLstml' exp. | FFA (n=12) | Category Pref, Non-pref | F(1,11)=15.22, **p=0.002** | F(1,11)=0.22, p=0.64 |
|  |  | Hemisphere R, L | F(1,11)=0.63, p=0.44 |  |
|  | PPA (n=14) | Category Pref, Non-pref | F(1,13)=34.92, **p=5\*10<sup>-5</sup>** | F(1,13)=5.43, **p=0.03** |
|  |  | Hemisphere R, L | F(1,13)=2.13, p=0.16 |  |
| Inverted 'Count20' exp. | FFA (n=9) | Category Pref, Non-pref | F(1,8)=13.89, **p=0.006** | F(1,8)=0.39, p=0.54 |
|  |  | Hemisphere R, L | F(1,8)=1.21, p=0.30 |  |
|  | PPA (n=10) | Category Pref, Non-pref | F(1,9)=4.43, p=0.06 | F(1,9)=0.005, p=0.94 |
|  |  | Hemisphere R, L | F(1,9)=9.05, **p=0.01** |  |

#### Control experiment 1: Upright ‘DBLstml’ experiment

To examine whether the effects we found in the ‘Count20’ experiment were task-dependent (Harel et al., 2014), we ran an additional control experiment (‘DBLstml’) with a new cohort of participants where we kept the same experimental paradigm but employed a different task (see Methods). We first replicated the expected category selectivity in the FFA and the PPA with higher activation to the central-preferred relative to the central non-preferred category (see Figure 5A and statistical results in Table 3). As in the ‘Count20’ experiment we also found here that for its preferred category PPA’s eccentricity-based modulations appeared more moderate that those found in the FFA (Figure 4). Specifically, normalized response analysis (see Methods and Figure 4B) showed that in FFA there was a decrease of ∼82% for faces from 0° to 8° (FFA-L: 81%±28%(SD), FFA-R: 84%±44%(SD); similar to ‘Count20’ exp. With ∼76% reduction) while in PPA there was a more moderate decrease of ∼13%±91%(SD) in PPA-L and ∼44%±47% in PPA-R for houses from 0° to 8° (cf. with PPA-L: ∼ 28%, PPA-R: ∼ 60% in the ‘Count20’ exp.).

Here, we found similar results to those found in the ‘Count20’ experiment. We measured these effects by computing the slopes of activation change with eccentricity in the parafovea (see Figure 4C and Table S4 at https://osf.io/m8czv/). For their preferred category both PPA and FFA showed negative slopes consistent with our results in the ‘Count20’ exp. (the BOLD eccentricity effect, one-sample, 2-sided Wilcoxon signed rank test: for houses in PPA (PPA-L: p=0.02, PPA-R: p=0.002) and for faces in FFA (FFA-L: p=0.0004, FFA-R: p=0.0001), see Table 3) with PPA showing quantitatively smaller slopes than those found in the FFA (2-way repeated-measures ANOVA with ROI _(FFA/PPA)_ X hemisphere _(R/L)_ on preferred category slope: effect of ROI (FFA/PPA): F(1,11)=16.79, p=0.002), Table 5). Importantly, for its non-preferred category PPA showed here too (as in the ‘Count20’ exp.) a positive slope (opposite to the expected BOLD eccentricity effect; one-sample 2-sided Wilcoxon signed rank test for faces: PPA-L (p=0.02), PPA-R (p=0.02)) that was qualitatively different than that found in the FFA for its non-preferred category (significantly negative slopes for houses: FFA-L (p=0.002), FFA-R (p=0.004), see Table 4).

In summary, here, with a different task and a new group of participants we found again results that are consistent with our original ‘Count20’ experiment.

#### Control experiment 2: Inverted ‘Count20’ experiment

In our behavioral study (Kreichman et al., 2020) we found that inversion exposed qualitatively different eccentricity-based perceptual modulation for houses than for faces. Therefore, here we were also interested to investigate whether this behavioral difference can be attributed to processing differences between FFA and PPA (i.e., whether stimulus inversion affects eccentricity-based modulations differently in FFA and PPA). Since upright and inverted stimuli were not presented in the same runs we were not able to directly compare their activation levels, and since inverted stimuli do not typically optimally activate these ROIs (e.g., (Yovel and Kanwisher, 2005; Gilaie-Dotan et al., 2010)) we did not calculate normalized responses.

In the FFA the results with the inverted stimuli for both preferred and non-preferred categories were similar (albeit noisier) to those with the upright stimuli, with negative slopes (i.e. BOLD eccentricity effect) for both categories (see Figure 5D-F, Table 3 and https://osf.io/m8czv/ Table S4; one-sample 2-sided Wilcoxon signed rank test: significantly negative slopes for inverted faces (FFA-L: p=0.003, FFA-R: p=0.001) and significantly negative slope for inverted houses (FFA-L: p=0.009, FFA-R: p=0.009)). We also found that the slopes were modulated by visual category such that eccentricity-based activation decline was faster for faces than for houses (see Table 6).

In the PPA the pattern found with inverted stimuli was different than that found with the upright ones. Specifically, for the inverted preferred category (inverted houses) the activation levels were mostly above baseline (see Figure 5F) but the slopes were very close to zero (see slope magnitude in Table S4 at https://osf.io/m8czv/, slope different from zero only at PPA-R (PPA-L: p=0.37, PPA-R: p=0.03, one-sample 2-sided Wilcoxon signed rank test), see Table 3). For the inverted non-preferred category (inverted faces) we found that the activations levels were around baseline, and importantly, there were no modulations of activity by eccentricity in both PPA-R and PPA-L (slopes not significantly different than zero: PPA-L (p=0.43), PPA-R (p=1); one-sample 2-sided Wilcoxon signed rank test, see Table 3).

In summary, for inverted stimuli the FFA showed an eccentricity effect for both categories while the PPA was much less affected by eccentricity for both categories.

Figure 6 presents a summary of the eccentricity-based reductions (i.e., slopes) for each of the ROIs according to their stimulus category preference for all the participants that participated in both the upright and the inverted ‘Count20’ experiments (n=11). As can be seen, while for the FFA a BOLD eccentricity effect was consistently found for preferred and non-preferred categories regardless of whether they were presented upright or inverted, for PPA there were qualitative changes of the eccentricity-based modulations (i.e., negative, zero and positive eccentricity-based modulations) found according to the visual category being viewed. While finding the anticipated BOLD eccentricity effect for its preferred category, we found no modulation by eccentricity to inverted faces, and a negative eccentricity effect for its non-preferred category of faces.

**Figure 6.**
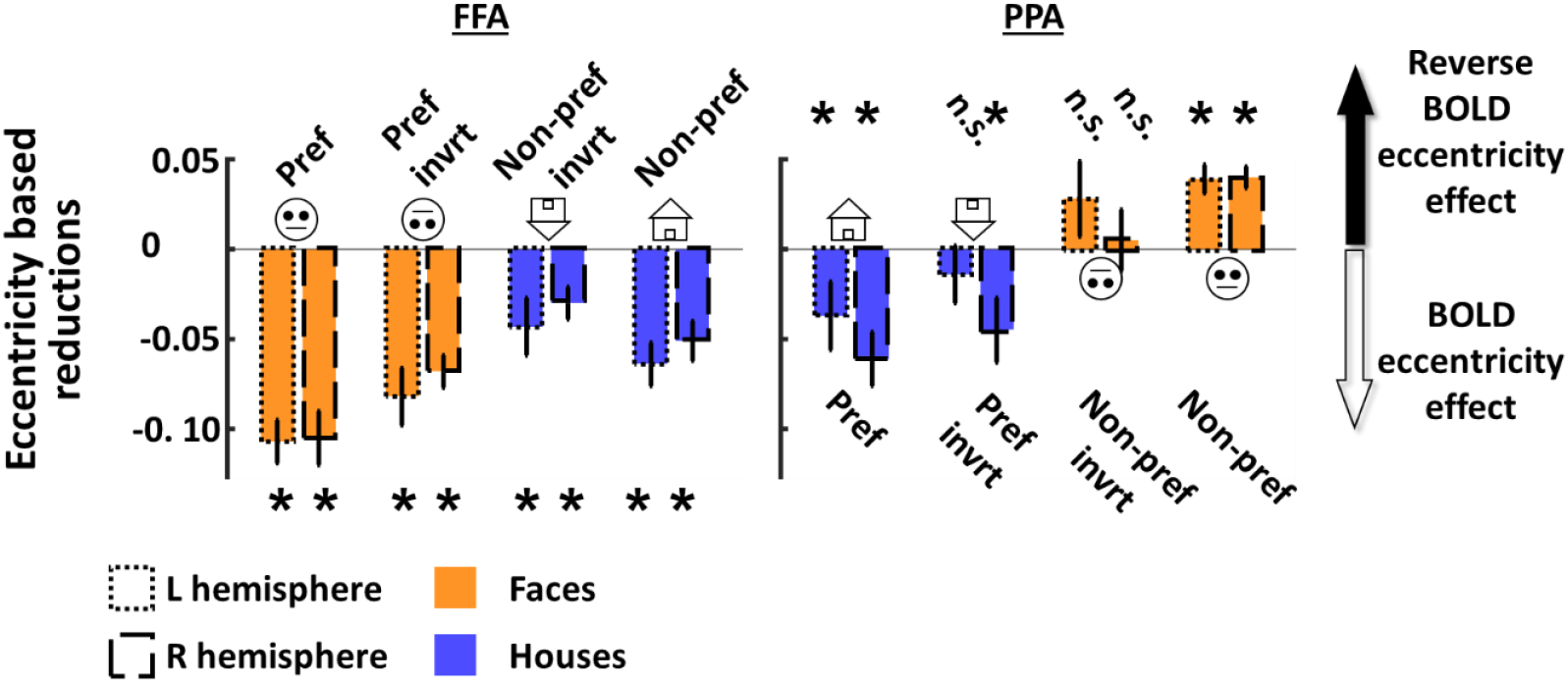
ROI eccentricity-based reductions (slopes) by category preference (n=11). For each region eccentricity-based reductions (y-axis, as measured by slopes) are presented by category preference (x-axis) from most preferred (left) to non-preferred (right). Faces in orange, houses in blue. Inverted categories are presented between these assuming inverted stimuli would be less preferred and less non-preferred as they may recruit non-dedicated networks. Negative slopes indicate a typical BOLD eccentricity effect, positive values a reverse (negative) BOLD eccentricity effect. Data of ‘Count20’ experiment presented for all n=11 participants with both upright and inverted data. Error bars represent SEM. Asterisks represent slopes significantly difference than 0.

### Category selectivity analysis

While category selectivity is often viewed as the “gold standard” for functionally partitioning high-level visual cortex into different category selective areas (Kanwisher et al., 1997; Epstein and Kanwisher, 1998; Saxe et al., 2006; Gilaie-Dotan et al., 2009, 2010; Grill-Spector et al., 2017b), this method is also criticized (e.g. for being condition/experiment dependent (Friston et al., 2006)). Furthermore, it is typically based on centrally presented stimuli. As FFA and PPA are two of the most investigated regions defined by functional localization criteria, here we were interested in also examining whether category selectivity in FFA and PPA are modulated by eccentricity (i.e., whether preferential category activations exist in parafoveal vision, see Figure 7).

**Figure 7.**
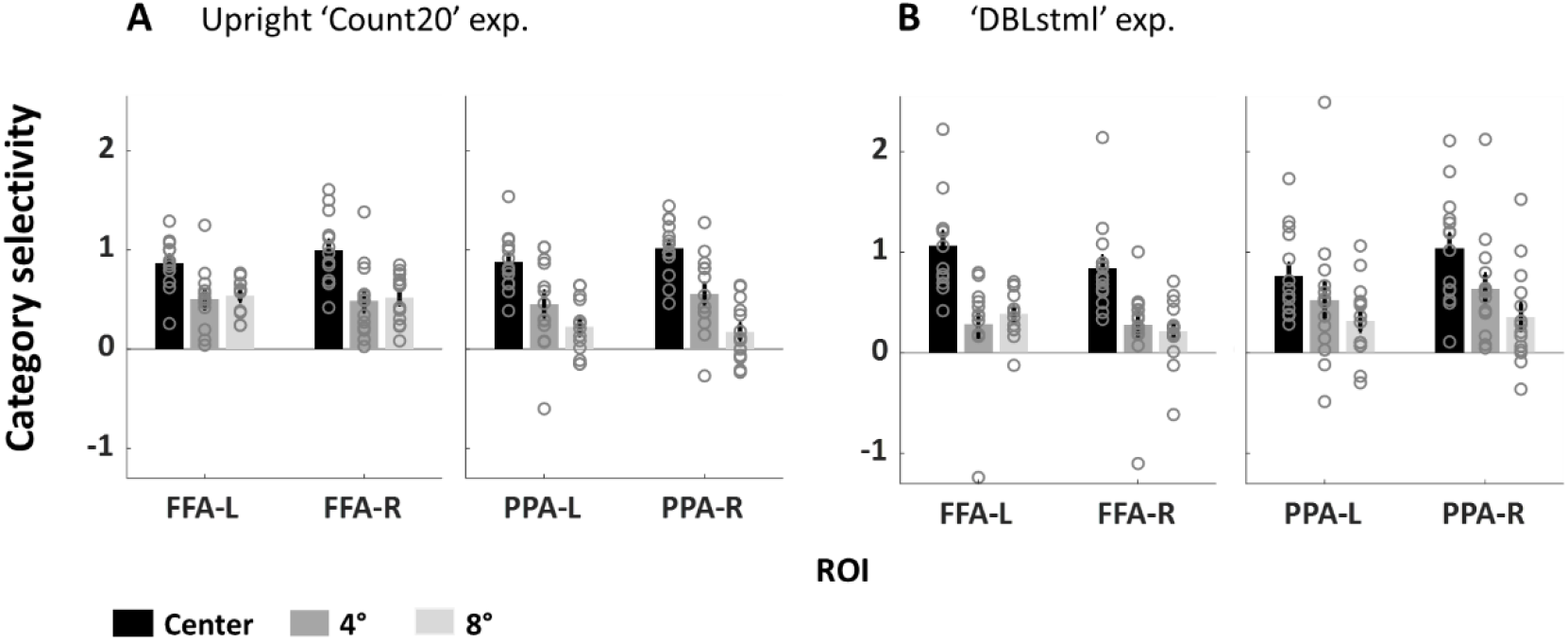
FFA and PPA category selectivity modulation by eccentricity. **(A)** Upright ‘Count20’ (n=14). **(B)** ‘DBLstml’ (n=14). Eccentricity demarcated by shades of black (central) to gray. In both experiments we found category selectivity was highest for central stimuli and decreased with eccentricity in all ROIs. However, in both FFA and PPA, category selectivity modulations in the parafoveal were not consistent across experiments. Error bars represent SEM.

Our main finding, which was consistent across experiments, was that there was a significant effect of eccentricity on category selectivity (1-way repeated measures ANOVAs on selectivity by eccentricity (0°/4°/8°) for each ROI: FFA-L: p=0.003, FFA-R: p=5*10^-5^, PPA-L: p=1*10^-5^, PPA-R: p=1*10^-7^) and this reflected highest category selectivity for central stimuli in both FFA and PPA (R and L) and significant reductions in category selectivity with eccentricity (see Figure 7 and Table 7). While the modulations of the selectivities within the parafovea (4° to 8°) were not consistent across our 2 experiments (see Figure 7), it is clear that across the 2 experiments for each of the ROIs the selectivity at 4° was not smaller than that found for 8° (see Figure 7, Table 7 and Table S4 at https://osf.io/m8czv/).

**Table 7.**
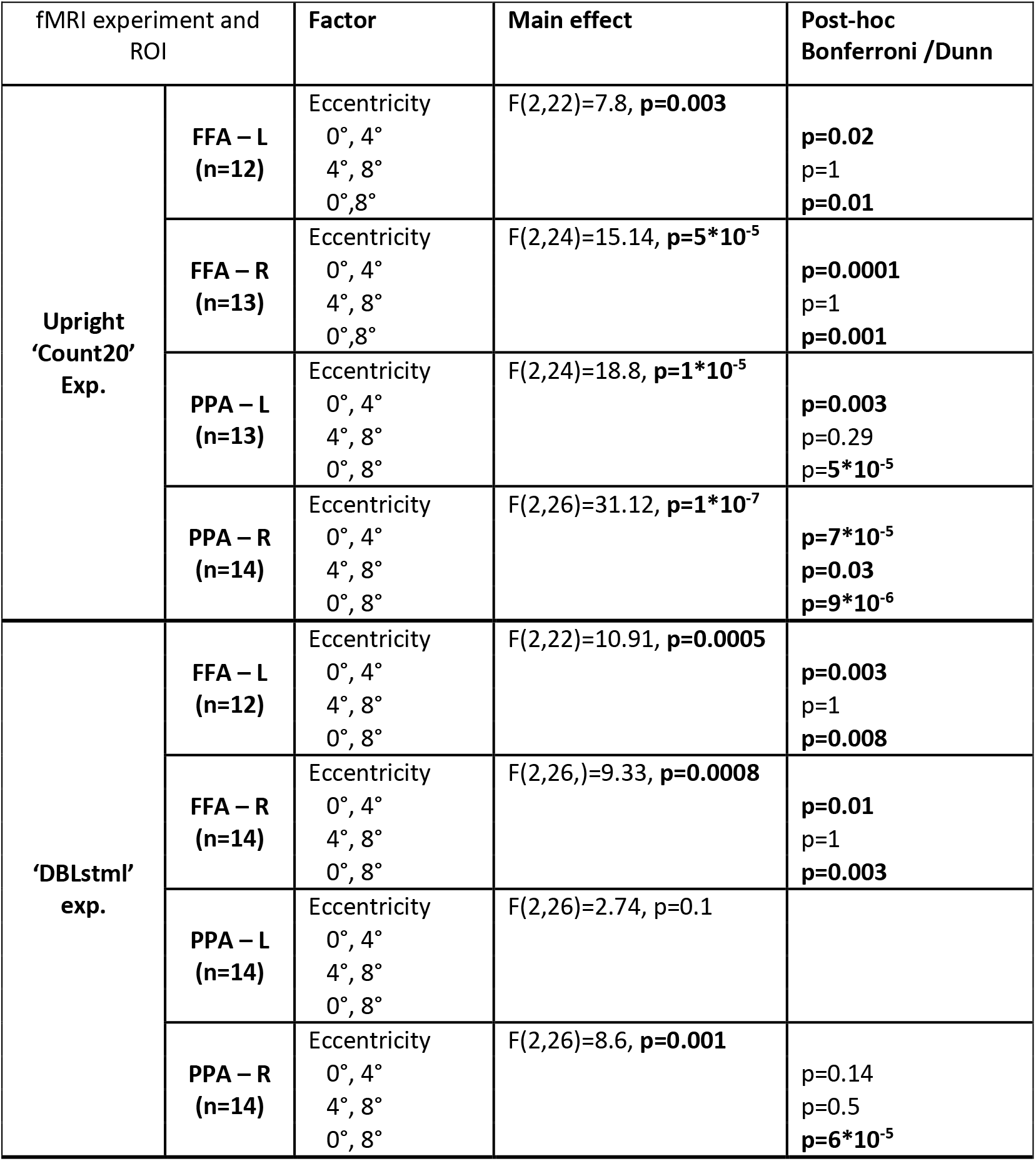
Eccentricity influence on category selectivity. For each region results of 1-way repeated measures ANOVA (with Eccentricity as factor) on category selectivity are presented (see Results). Significant effects in bold. Note that overall there were significant influences of eccentricity on category selectivity (highest at the center, see Figure 7).

## Discussion

Here we investigated how eccentricity modulates activations in FFA and PPA, face- and place-selective regions in human ventral visual cortex, given our recent behavioral findings indicating that perception for faces declines faster with eccentricity than for houses (Kreichman et al., 2020). We hypothesized that (i) FFA will be more sensitive to eccentricity than PPA given its foveal bias and PPA’s peripheral bias (Levy et al., 2001), and that (ii) each region will show a characteristic region-specific BOLD eccentricity effect mirroring the behavioral eccentricity effect (Carrasco et al., 1995; Wolfe et al., 1998) and will be independent of the viewed category. In a set of fMRI experiments we found, in line with our hypothesis, that FFA showed bigger BOLD eccentricity effects in the parafovea (≤8°) than those found in PPA. However, within each region we found that the reduction rate was modulated by visual category, and importantly that the effect of category on PPA’s modulation was more profound (qualitative) than that found for FFA (showed only quantitative changes in the BOLD eccentricity effect). Specifically, we found that PPA’s activity was modulated by eccentricity from reductions to enhancements depending on the viewed category. In addition, both regions’ characteristic category-selectivity reduced in the parafovea.

### BOLD eccentricity effect modulated by visual category in FFA and PPA

FFA showed a more rapid decline of activation with eccentricity than PPA did, even when considering the category-related differences, and this was in line with our initial hypothesis. This finding is line with our behavior results that led to this study (Kreichman et al., 2020) where we found that face discrimination performance decreased more rapidly than house perception (at ≤4°). Regardless of these different sensitivities to eccentricity, for each of these regions we found, in contrast to our assumption, that their eccentricity-based activation modulations were modulated by visual category. This finding challenges our hypothesis for several reasons. Firstly, processing within each region may be region-specific rather than category-specific (i.e. display similar computational properties across different visual categories (Gilaie-Dotan et al., 2008)). Furthermore, neuroanatomical cytoarchitectural investigations reveal different compartments within ventral cortex [(Weiner et al., 2017) FG1-FG4, likely corresponding to face- (FG4 to mFus-faces likely corresponding to our FFA ROI) and place- (FG3 to CoS-places likely corresponding to our PPA ROI) sensitive regions]. Such cytoarchitectural characteristics may suggest that the computation each region is performing is hardwired and not category-specific. While our results that the BOLD eccentricity effect is modulated by the viewed category may be surprising, they could be in line with the findings that processing in ventral cortex may reflect interactions between different properties (e.g., disparity and shape (Gilaie-Dotan et al., 2002) or disparity, shape, and viewpoint (Janssen et al., 2000)). This may suggest that computational processes within these ventral regions are sensitive to eccentricity on top of their category preference as we have found in the PPA and to a much lesser extent in the FFA. While in the FFA all categories undergo activation reductions with eccentricity (i.e., negative slopes for all visual categories reflecting a BOLD eccentricity effect), in the PPA the activation changes by eccentricity are profoundly affected by the visual category – from reduction in activation with growing eccentricity (a BOLD eccentricity effect for houses) to enhancement in activation with eccentricity (reverse BOLD eccentricity effect for faces). This means that PPA undergoes *qualitatively* different eccentricity-based modulations by the viewed category. Furthermore, these results may support the assumption that representations in ventral visual cortex are distributed such that multiple regions are responsive to multiple categories (e.g. (Ishai et al., 1999)).

### Investigating peripheral processes in high-order visual cortex with and without accounting for CMF

Some previous studies investigating parafoveal processing in high-order visual cortex compensated for the CMF (Levy et al., 2001, 2004; Malach et al., 2002; Schwarzlose et al., 2008) finding that FFA shows a significant foveal bias and PPA shows a strong peripheral bias. However, it is unclear how activation levels are modulated when the stimulus keeps its world size and thus its retinal size. Furthermore, some of the peripheral stimuli employed in these studies included multiple peripheral elements vs a single foveal element potentially confounding the visual field findings. Here to parametrically investigate how eccentricity influences visual activation we kept stimulus size constant. We deliberately chose this to mimic the constancy in stimulus retinal size following a shift of gaze across the visual field. Our design enabled us to examine more closely modulations within the parafovea and importantly examine if they are region-specific (as we anticipated) or sensitive to the viewed category as well as to inversion. A study focusing on FFA while not accounting for CMF and using upright face stimuli (Yue et al. 2011) found a significant BOLD eccentricity effect (i.e., activation reduction as a function of distance from central vision), which is in line with our findings in the FFA for faces. Interestingly, while we found eccentricity-based enhancement of activation in PPA for faces (i.e., a reverse BOLD eccentricity effect), they find no eccentricity-based modulations in the PPA for faces, and their result may also be considered as a qualitative deviation from an anticipated BOLD eccentricity effect.

### Limitations

While our results clearly indicate that eccentricity in the parafovea modulates activation levels in FFA to stronger extent than in PPA, and that within each of the regions activation modulation is category-dependent, there are certain limitations of our study that must be taken into account. The fact that we did not directly map pRFs (Dumoulin and Wandell, 2008; Kay et al., 2015; Silson et al., 2015; Finzi et al., 2021) prevents us from determining whether the sensitivity of each region is uniform or varies according to visual field sensitivities. FFA has shown consistent preference to central vs. peripheral vision (Levy et al., 2001; Hasson et al., 2002). In addition, most population receptive fields (pRFs) in FFA cover the fovea for varying pRF sizes (Kay et al., 2015; Grill-Spector et al., 2017a; Gomez et al., 2018), and thus while foveal stimuli are likely to activate most of the FFA, stimuli at eccentricities beyond central vision (in our case the 4° and 8° conditions) are likely to activate a smaller part of the FFA. Therefore, the observed rapid reduction in FFA’s BOLD magnitude with eccentricity may have resulted from reduced activation in the activated voxels and/or from a reduction in the number of activated voxels. PPA on the other hand is more sensitive to visual field organization (Silson et al., 2015, 2016) with a more retinotopic-like representation where pRFs seem to be centered across different eccentricities in the visual field.

Therefore, in the PPA we may be sampling different groups of neurons when we compare activations to the different eccentricities. Furthermore, the fact that we have not employed fMRI adaptation methods (Grill-Spector and Malach, 2001; Gilaie-Dotan and Malach, 2007; Gilaie-Dotan et al., 2008, 2010) prevents us from precisely determining whether the category-based reductions within each region reflect modulations within the same population of neurons or whether they reflect within-voxel modulations of different sub-populations of neurons. In addition, our stimuli were not optimal for activating PPA. This is due to the fact that our localizer was limited to ∼6°x6° and therefore our analyses may have not captured PPA’s sensitivity to eccentricities beyond that, and our stimuli (sized ∼2°x2.5°) may have been too small to address PPA’s peripheral bias (Levy et al., 2004). Therefore, it may be the case that more specific investigations of PPA across the visual field may allow more precise characterization of its functional sensitivity relative to those evident in our study. It is also unclear whether FFA and PPA will show BOLD eccentricity effects for categories not tested in our study and whether these would be within the range found here. Furthermore, our study focused on FFA and PPA and it is unclear whether activity in additional category-sensitive regions within high-order visual cortex are also sensitive to eccentricity across the visual field.

### Summary

While further investigations are required to address additional aspects not investigated in our study, our results highlight important properties of high-level visual cortex. As perception changes with growing distance from central visual field (e.g., Kreichman et al., 2020; Akselevich and Gilaie-Dotan, 2022), here we found that two of the most prominent regions in ventral high-order visual cortex (FFA and PPA) show significant eccentricity-based modulations of BOLD activity within the parafovea (a BOLD eccentricity effect). Importantly, these modulations are not only driven by eccentricity but also by the viewed category. In FFA these BOLD modulations were quantitative such that for all investigated categories a BOLD eccentricity effect was found. For PPA on the other hand, modulations were much more substantial with qualitative modulations by the viewed category, from a BOLD eccentricity effect as we anticipated, to an unexpected reverse BOLD eccentricity effect. We propose that PPA’s activity may reflect not only a peripheral bias, but also antagonism to central visual field processes (such as those that are driven by central upright faces but not by central inverted faces that elicit activation beyond central-vision-related networks (Gilaie-Dotan et al., 2010)).

## Acknowledgments

We thank Rafi Malach and Michal Harel for their support and Kalanit Grill-Spector for her comments and suggestions. The study was funded by ISF grant 1485/18 to SGD and Council for Higher Education Levtzion scholarship to OK.

